# The nuclear receptor ROR-alpha is a critical myogenic regulator of the cardiomyocyte transcriptome, including the alpha-1A adrenergic receptor

**DOI:** 10.64898/2026.01.11.698261

**Authors:** RS Nagalingam, JY Beak, LG Kirkland, RL Lutze, W Huang, D Akkina, M Goyal, C Kang, A Zhao, B Alcira, A Aghajanian, K Gerrish, HS Kang, AM Jetten, BC Jensen

## Abstract

**Aims:** We recently found that the nuclear receptor retinoic acid-related orphan nuclear receptor alpha (RORα) protects against angiotensin II-induced cardiac hypertrophy and promotes cardiomyocyte mitophagy. The underlying molecular basis for these salutary effects remains unclear.

**Methods:** We used RNA microarrays to profile the cardiac transcriptomes of “staggerer” (RORα^sg/sg^) mice that carry a naturally occurring mutation in the ligand-binding domain of RORα, resulting in a global loss-of-function genetic model. We then used genetic and pharmacologic loss-and-gain of function studies in cultured cardiomyocytes to ascertain whether RORα regulates transcription of *Adra1a*, the gene that encodes the alpha-1A-adrenergic receptor (α1A-AR).

**Results:** The absence of functional RORα results in broad transcriptional changes in the heart providing a likely molecular basis for the RORα^sg/sg^ cardiac phenotype. *In vivo* and *in vitro* studies confirmed that RORα directly regulates *Adra1a* transcription. This effect is enhanced by hypoxia.

**Conclusions:** Collectively these findings position RORα as a previously unrecognized central regulator of the cardiac myogenic transcriptome and the first recognized transcriptional regulator of *Adra1a* in cardiomyocytes. Future studies will probe the contribution of RORα-mediated transcriptional regulation of *Adra1a* to both the response to cardiomyocyte injury and maintenance of circadian biology.

## Introduction

The progression from index cardiac injury to clinically evident heart failure (HF) is governed in part by the transcriptional programs that collectively regulate critical biological processes including contractile protein isoform expression, calcium handling, and cardiomyocyte metabolism[1]. The importance of the nuclear receptor superfamily of transcription factors in coordinating these programs is increasingly recognized[2]. Nuclear receptors are ligand-activated transcription factors that include receptors for lipid-soluble compounds like steroid hormones, vitamins, and cholesterol metabolites. The ligands for many of these receptors were not known at the time of their naming, leading to their initial categorization as “orphan receptors”. Currently, however, many nuclear receptors with recognized ligands are targets of either approved or experimental drugs for a wide and expanding range of diseases[3]. Indeed, mineralocorticoid receptor antagonists prolong survival in patients with all stages of HF[4], demonstrating the therapeutic potential for targeting nuclear receptors as much-needed novel treatments for this devastating clinical syndrome.

The retinoic acid-related orphan nuclear receptor (ROR) subfamily consists of three members, RORα, RORβ, and RORγ (NR1F1-3), with tissue-specific and context-dependent expression. Each ROR exhibits a typical nuclear receptor domain structure, consisting of an N-terminal domain, a highly conserved DNA-binding domain (DBD) with two C2-C2 zinc finger motifs, a ligand-binding domain (LBD), and a hinge domain between the LBD and DBD. The RORs regulate transcription by binding to ROR response elements (ROREs) in regulatory regions of target genes that consist of an RGGTCA consensus preceded by an A/T-rich sequence[5]. All three RORs play crucial roles in lipid metabolism, immune cell development, and regulating circadian rhythm[6]. However, there was no recognized function for RORs in the heart prior to two reports showing that RORα mitigates myocardial ischemia/reperfusion injury and diabetic cardiomyopathy[7, 8].

We initially investigated the effects of RORα in the heart using the well-established “staggerer” (RORα^sg/sg^) mice that globally lack functional RORα[9]. RORα^sg/sg^ mice, studied for over 60 years as a model of cerebellar dysfunction, carry a naturally occurring deletion in *NR1F1* (the gene that encodes RORα) that prevents the translation of the ligand binding domain of RORα[9]. These mice exhibit an ataxic gait[10], tremor, and small body size with reduced adiposity[11], enhanced insulin sensitivity[12], and mildly reduced blood pressure[13]. We found that RORα^sg/sg^ mice develop exaggerated myocardial hypertrophy and contractile dysfunction after 14 days of angiotensin II exposure and that RORα is downregulated in the setting of mouse and human HF[14]. We then extended those findings to show that RORα adaptively regulates cardiomyocyte mitophagy to enhance mitochondrial function in the basal state and after injury[15]. Taken together, these findings suggest that RORα may play protective roles in HF. The mechanisms underlying these putative cardioprotective benefits and the functions of RORα in uninjured cardiomyocytes remain uncertain.

In this manuscript, we probed for potential underlying mechanisms by carrying out RNA microarray on the hearts of RORα^sg/sg^ mice and wild type controls. We found broad and high magnitude alterations in the abundance of thousands of transcripts, including those in key pathways related to sarcomeric structure and function and cardiac fibrosis. Using this unbiased approach we identified markedly lower abundance of *Adra1a*, the gene that encodes the alpha-1A adrenergic receptor (α1A-AR), in RORα^sg/sg^ hearts. Given the longstanding interest of our lab in the α1A-AR, we conducted focused experiments to demonstrate direct regulation of *Adra1a* transcription by RORα.

## Methods

### Experimental animals

Heterozygous “staggerer” mice on a C57BL/6J background were purchased from Jackson lab and maintained as previously described[16]. Homozygous male staggerer mice (RORα^sg/sg^), the products of heterozygous breeding, and wild type littermates were used in all experiments at 12-16 weeks of age. Homozygous cardiomyocyte-specific RORα knockout mice (RORα CMKO) were generated by crossing floxed RORα mice with LoxP sites flanking exons 9-11 (Anton Jetten, NIEHS, Research Triangle Park, NC) with αMHC-Cre mice (Dale Abel, University of Iowa). All animal studies followed the NIH Guide for the Care and Use of Laboratory Animals and animal protocols were approved by the NIEHS Animal Care and Use Committee and the University of North Carolina Institutional Animal Care and Use Committee. For euthanasia, animals were anesthetized with 4% isoflurane by continuous inhalation until the toe pinch reflex was absent then underwent cervical dislocation.

### Microarrays

Total RNA was isolated from hearts of 12–16-week-old WT (n=5) or RORα^sg/sg^ (n=4) mice using Invitrogen PureLink RNA mini-kit (ThermoFisher). Microarray analysis was carried out by the NIEHS Molecular Genomics Core using Agilent (Palo Alto, CA) whole genome mouse oligo arrays[17]. Data obtained with the Agilent Feature Extraction software (v7.1) were further analyzed with OmicSoft Array Studio (v6.0) software. Gene expression analysis and heatmap generation were conducted as previously described[18]. All the data on microarray have been deposited in the NCBI’s Gene Expression Omnibus number under GSE98892.

### Quantitative real-time reverse transcriptase PCR (qRT-PCR)

Total RNA from WT and RORα^sg/sg^ mouse hearts was isolated with an RNeasy mini kit (Qiagen, Valencia, CA) or RNAqueous^®^ Micro RNA isolation kit (Ambion, Austin, TX), respectively, following the manufacturer’s instructions. RNA was reverse-transcribed using a High-Capacity cDNA Archive Kit (Applied Biosystems, Foster City, CA). qRT-PCR reactions were carried out in triplicate in a LightCycler® 480 System (Roche, Indianapolis, IN, USA) using either probe/primer sets or SYBR Green I. Relative quantitation of PCR products used the ΔΔCt method relative to two validated reference genes (*Tbp* and *Polr2a*). All probes and primers were from Roche or ThermoFisher.

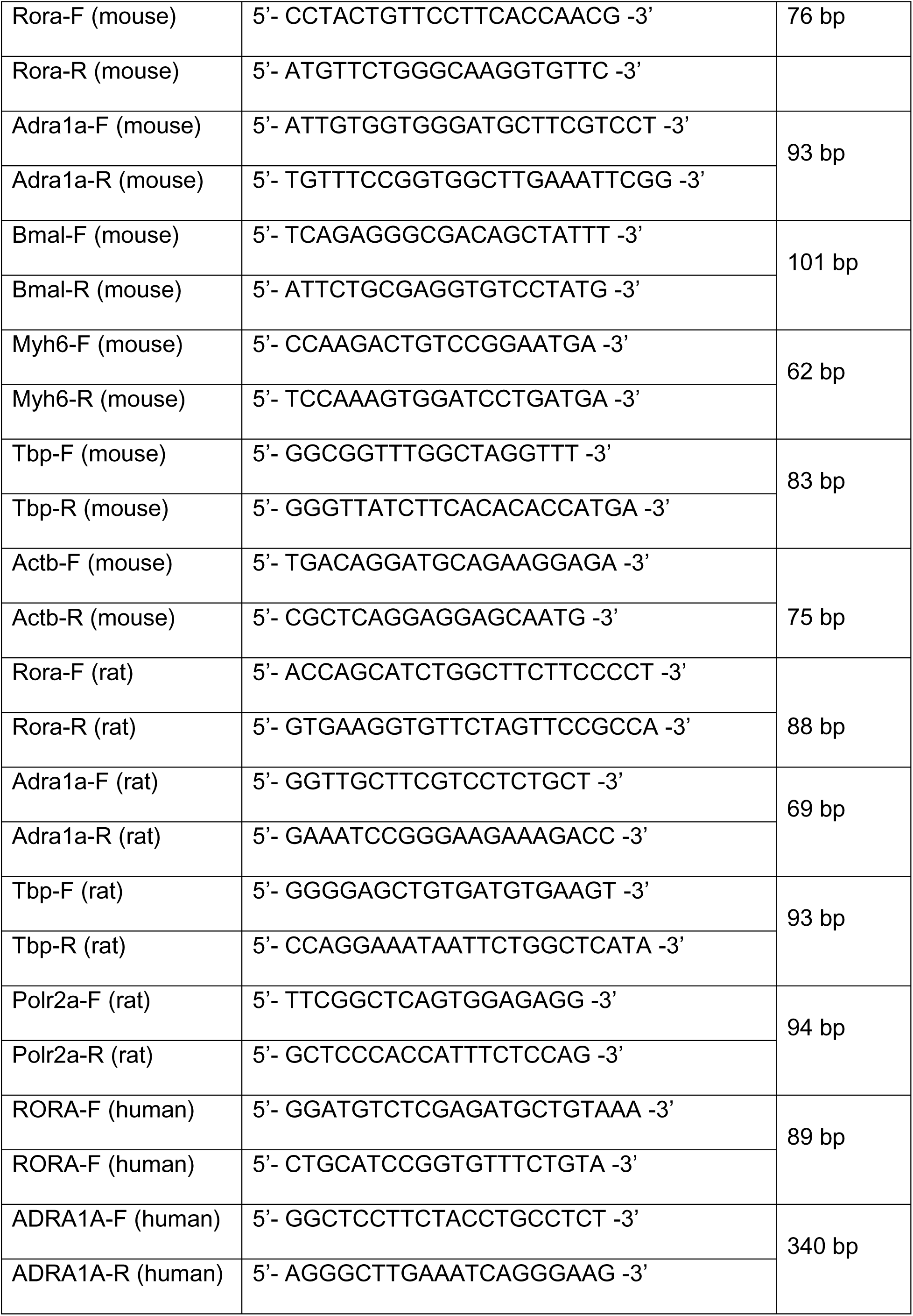
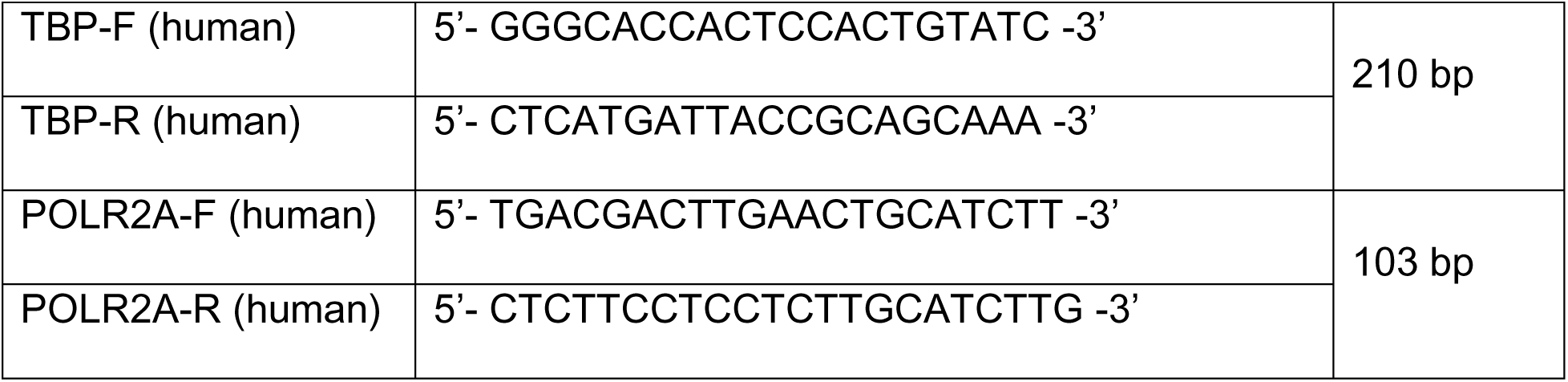

### Hypoxia chamber

Mice were housed in a hypoxia chamber (Oxycycler) with internal dimension (76x51x51 cm) sufficient to hold 4 cages with 4 mice per cage as previously described[19]. Inflow rate (nitrogen mixed with oxygen or room air) was ∼3.1 ft^3^/hr and 4 holes (0.7 cm diameter) were opened at the bottom of each of the chamber’s 3 sides. Drierite (#21909-5000, Acros, Fair Lawn, NJ) and soda lime (#36596, Alfa Aesar, Haverhill, MA) were placed at the bottom of the chamber in 12x10x5 cm trays (∼260g and 200g, respectively). CO_2_ was measured using a Cozir Wide-Range sensor and GasLab software, (#CM-0123 http://CO2meter.com, Ormond Beach, Fl). The hypoxia chambers have internal fans to prevent gradients in composition of the atmospheres. CO_2_ was automatically recorded at 5 min intervals. FIO_2_ was lowered from 21% to 10% (hypoxia) in ∼3 h. Control (21% O_2_, normoxia) mice were maintained in an identical chamber. Hypoxia chambers were generously provided by Dr. James Faber.

### Neonatal rat ventricular myocyte (NRVM) cultures, immunocytochemistry and lentiviral infections

Female Sprague-Dawley rats and newborn litters were from Charles River Laboratories (Wilmington, MA, USA). Neonates were deeply anesthetized with 4% inhaled isoflurane then euthanized by decapitation and NRVMs were isolated as previously described[20]. Experiments were carried out after 36-48 hours of serum starvation in the presence of insulin, transferrin, and BrdU. NRVM hypoxia was induced by incubation in a 37°C hypoxia chamber with 1% O_2_ after infection with lentivirus containing either scrambled short hairpin RNA (shControl) or shRNA specifically targeting rat RORα (iO51217 or iV051217, Abm, Richmond, BC, Canada). shRORα target sequences:

TGTCATTACGTGTGAAGGCTGCAAGGGCT, ACCTACAACATCTCAGCCAATGGGCTGAC, GGACTGGACATCAATGGGATCAAACCCGA, AGAGGTGATGTGGCAGTTGTGTGCTATCA

### Culturing and maintenance of human induced pluripotent stem cells (hiPSCs)

Healthy control human iPSC Lines (Female, SCTi003-A and Male, SCTi004-A) were obtained from StemCell Technologies (Canada). Cryopreserved hiPSCs were thawed and plated onto 2% Matrigel-coated (354230, Corning, USA) 6 well plate in mTeSR plus medium (100-0276, StemCell Technologies, Canada) supplemented with 5µM Y-27632 (72304, StemCell Technologies, Canada) and 10% cloneR2 (100-0691, StemCell Technologies, Canada) to enhance the post thaw survival. After 24 hours, the culture medium was replaced with fresh mTeSR plus medium, which was subsequently changed daily. When cultures reached approximately 70–80% confluency, hiPSCs were passaged using ReLeSR (00-0483, StemCell Technologies, Canada) and replated on a Matrigel-coated 6 well plates with the seeding density of 2 x 10^5^ cells per well. hiPSCs were monitored routinely for pluripotent colony morphology and the absence of spontaneous differentiation.

### Differentiation of hiPSC-derived cardiomyocytes

Differentiation of hiPSCs into cardiomyocytes was performed following a small molecule-based protocol adapted from Burridge et al[21]. Briefly, hiPSCs were seeded on Matrigel-coated plates and cultured until colonies reached ∼80% of confluency. On day 0, differentiation was initiated by replacing the medium with RPMI 1640 containing 2% B27 supplement minus insulin (A1895601, ThermoFisher Scientific, USA), 60 ng/µL Activin A (120-14P-50UG, PeproTech, USA), 12 µM CHIR-99021 (100-1042, StemCell Technologies, Canada), and 50 µg/mL L-ascorbic acid (A5960, Sigma-Aldrich, USA). On days 1 and 2, cells were fed daily with fresh RPMI 1640 supplemented with 2% B27 supplement minus insulin, 5 µM IWR-1 (72562, StemCell Technologies, Canada), and 50 µg/mL L-ascorbic acid. On day 4, the medium was replaced with RPMI 1640 containing 2% B27 supplement minus insulin and 5 µM IWR-1. From day 6 onward, cells were maintained in RPMI 1640 with 2% B27 Supplement with insulin (17504044, ThermoFisher Scientific, USA), refreshed every other day. Spontaneous contractile activity was typically observed between days 7–9 of differentiation.

### Treatment with RORα inverse agonist (SR3335) and agonist (SR1078)

NRVMs were treated with vehicle or RORα inverse agonist, SR3335 20μM[22] or agonist, SR1078 2μM[23] for 6 hr and then incubated in a hypoxia chamber (1% O_2_) after pretreatment with BFA (100nM) for 2 hr.

### Chromatin immunoprecipitation-PCR

To demonstrate transcriptional regulation of *Adra1a* by RORα, H9c2s were washed in PBS containing protease and phosphatase inhibitor cocktail (PIC; ThermoFisher) and then fixed in 1% formaldehyde at 25°C for 8 min for cross-linking and followed by quenching by 0.125 M glycine at 25°C for 8 min as previously described[14]. The amount of chromatin immunoprecipitated DNA (ChIP’ed-DNA) relative to each input DNA was determined by quantitative PCR in triplicate with primers amplifying *Adra1a* or a nontargeting sequence.

Primers used for qPCR following chromatin immunoprecipitation assay:

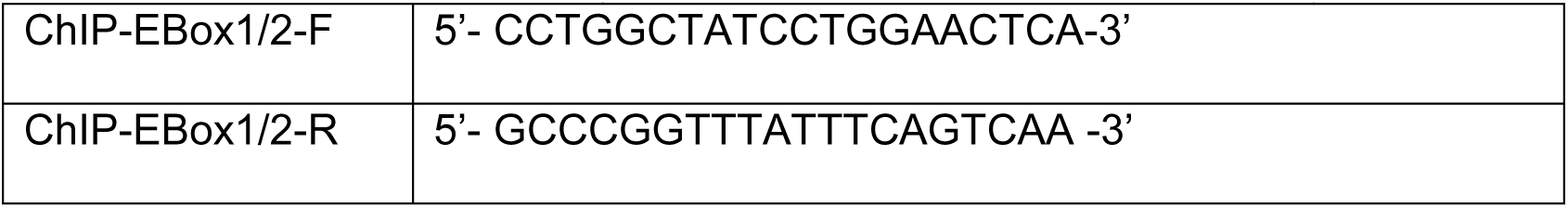

### Luciferase reporter assay

The luciferase reporter construct was generated by cloning an ∼819 bp proximal promoter region upstream of the mouse *Adra1a* gene into the pGL4.10 vector, yielding the pGL4.10-ADRA1α-Pro construct. A mutant version (M1-pGL4.10-ADRA1α-Pro) was created by site-directed mutagenesis, in which the RORE(2) within the *Adra1a* proximal promoter was altered from TGGGTCA to TG<u>A</u>G<u>C</u>CA. Both wild-type and mutant constructs were synthesized by GenScript. HEK293 or H9C2 cells were co-transfected with either pGL4.10-ADRA1α-Pro or M1-pGL4.10-ADRA1α-Pro together with an RORα expression vector (p3xFLAG-CMV-RORα) or an empty vector control (p3xFLAG-CMV). Transfections were performed under low-serum conditions using Lipofectamine 3000 (18324012, ThermoFisher Scientific) for 24 h. To normalize for transfection efficiency, cells were co-transfected with either a Renilla luciferase vector (pRL) or β-galactosidase (β-Gal). Twenty-four hours after transfection, cells were lysed with Passive Lysis Buffer and luciferase activity was quantified using the Dual-Luciferase Reporter Assay System (E1910, Promega) on a CLARIOstar multimode plate reader. β-Gal activity was measured using the β-Galactosidase Detection Kit II (Takara) following the manufacturer’s instructions. All transfections were performed in triplicate, and each experiment was independently repeated at least three times.

### Western blotting

Cell lysates were prepared in RIPA lysis buffer supplemented with PhosSTOP (Roche Diagnostics) and protease inhibitor cocktail (Roche Diagnostics). Subsequently, 30mg of protein samples were incubated in 4% SDS sample buffer, including 2% β-mercaptoethanol, for 10 min at 70°C. Then protein samples were resolved in 4-20% criterion TGX Stain-free precast gels (Bio-Rad) and transferred to PVDF membranes by using Transblot turbo transfer system (Bio-Rad). Membranes were blocked in 5% milk-Tris-buffered saline-Tween 20 and incubated in primary antibody overnight at 4°C and then secondary horseradish peroxidase (HRP)-conjugated antibodies for 1 h at room temperature. Images were generated using Clarity ECL western blotting substrate (Bio-Rad) and the images were captured using a ChemiDoc Imaging System (Bio-Rad). Protein quantification was conducted using Image J (NIH, USA).

#### Antibodies

Total ERK1/2 (#9102, 1:1000) and phospho-ERK1/2 (#4370, 1:2000) were from Cell Signaling. GAPDH (60004-1-Ig, 1:10,000) was from Proteintech (Rosemont, IL).

### Statistics

All results are presented as mean ± SEM. Comparisons were made using t-test (groups of 2) or one-way ANOVA (groups of 3) with Tukey’s post-hoc analysis (GraphPad Prism).

## Results

### The absence of functional RORα is associated with broad transcriptomic changes in RORα^sg/sg^ hearts

Our published characterization of the RORα^sg/sg^ (“staggerer”) revealed that global loss of RORα function is associated with a small, hypocontractile, and mildly fibrotic heart[14]. The basis for this pathological basal cardiac phenotype has not been established.

To investigate the molecular targets and pathways regulated by RORα in the heart, we performed an unbiased microarray based unbiased transcriptomic analysis in wild-type (WT) and RORα^sg/sg^ mouse hearts harvested at synchronized timepoints to negate potential differences induced by the fact that RORα is a known peripheral clock gene (**Figure 1A**). Principal component analysis (PCA) revealed a clear segregation between groups of transcriptomic profiles between WT (n=5) and RORα^sg/sg^ (n=4) hearts (**Figure 1B**). Using a Benjamini-Hochberg False Discovery Rate of <0.05 as a cutoff, we identified 9653 transcripts that were differentially expressed in RORα^sg/sg^ and WT hearts. Of these transcripts, 4418 were >2.0-fold more abundant and 3547 were >2.0-fold less abundant in RORα^sg/sg^ compared with WT hearts (**Figure 1C**) with relatively modest inter-replicate variation in transcriptomic profile (**Figure 1D**).

**Figure 1.**
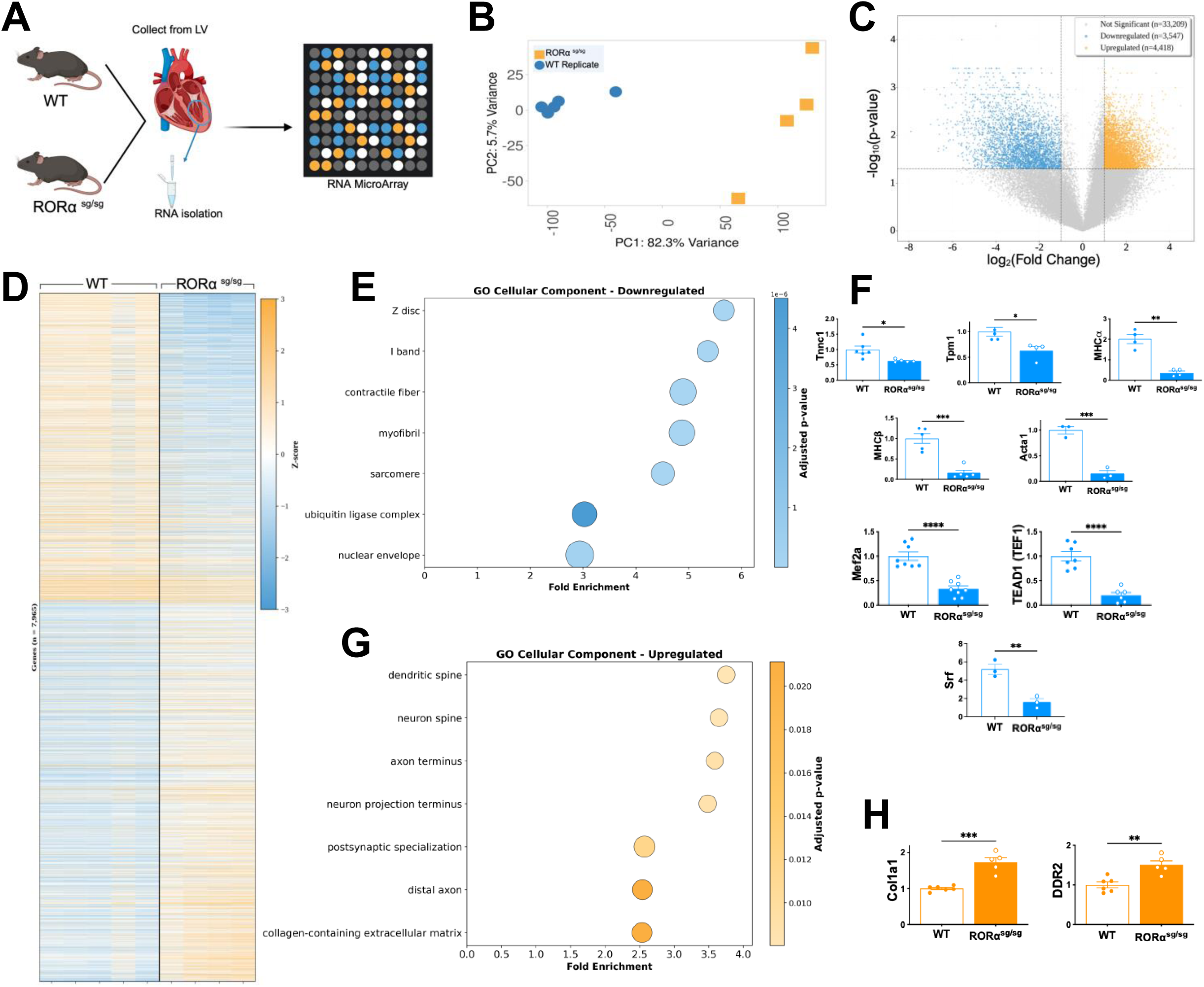
Transcriptomic analysis of the “staggerer” (RORa^sg/sg^) mouse heart, a genetic loss-of-RORa function model. (**A**) Experimental design schematic; **(B)** Principal component analysis (PCA) of WT and RORα^sg/sg^ mouse heart replicates; **(C)** Volcano plot depicting differentially expressed genes in RORα^sg/sg^ and WT hearts, with significantly upregulated and downregulated transcripts; (**D**) Hierarchical clustering heatmap illustrating distinct transcriptional profiles between WT and RORα^sg/sg^ hearts; **(E)** Gene Ontology (GO) Cellular Component term for 2-fold downregulated transcripts; (**F**) Quantitative RT-PCR of selected sarcomeric genes and transcription factors; (**G**) GO Cellular Component term for 2-fold upregulated transcripts KEGG pathway map of circadian entrainment highlighting altered components (red boxes) in RORα^sg/sg^ hearts; (**H**) Quantitative RT-PCR of selected fibrosis-related genes *p<0.05, **p<0.01, ***p<0.001, ***p<0.0001 vs WT.

We then carried out functional enrichment analysis of these two-fold regulated transcripts using two established platforms to probe for pathways that might be particularly affected by the loss of RORα function. In the set of downregulated transcripts, multiple myofibrillar Gene Ontology (GO) Cellular Component terms (Z-disc, I band, contractile fiber, myofibril, sarcomere) were the most strongly enriched (**Figure 1E**). Quantitative reverse transcription PCR (qRT-PCR) confirmed that the abundance of selected canonical sarcomeric genes (*Tnnc1, Tpm1, Myh7, Myh6, and Acta1*) was markedly lower in RORα^sg/sg^ than WT hearts (**Figure 1F**), consistent with our published atrophic and hypocontractile phenotype[14]. Interestingly, we also found that the abundance of 3 critical myogenic transcription factors (*Mef2a, Tead1, Srf*) was more than 2-fold lower in RORα^sg/sg^ mouse hearts (**Figure 1F**). This intriguing finding suggested the potential that RORα might function in a meta-regulatory transcriptional myogenic role in the heart, contributing to the extremely broad transcriptomic differences in our microarray dataset (**Figure 1F**).

GO Cellular Component analysis of the transcripts 2-fold more abundant in RORα^sg/sg^ than WT hearts revealed fewer statistically significant terms (**Figure 1G**). Confirmatory qRT-PCR identified upregulation of two canonical transcripts from the collagen-containing extracellular matrix term (*Col1a1* and *DDR2*) providing some transcriptional corroboration for the increased fibrotic burden in RORα^sg/sg^ hearts [14]. KEGG functional analysis revealed enrichment in transcripts related to cardiomyopathies as well as autophagy (**Supplemental Figure S1A**)—particularly interesting to us given our previous finding that RORα regulates cardiomyocyte mitophagy [15]. KEGG analysis of upregulated pathways identified fewer statistically significant terms, consistent with our GO analysis (**Supplemental Figure S1B**).

Collectively, these findings are consistent with the concept that our published RORα^sg/sg^ cardiac phenotype of impaired developmental cardiomyocyte hypertrophy and fibrosis arises from aberrant transcriptional regulation due to the absence of RORα. The breadth and magnitude of the observed changes also suggest that RORα plays a larger role in regulation of cardiac transcription than previously recognized.

### Loss of ROR⍺ function diminishes the abundance of *Adra1a*, the transcript encoding the α1A adrenergic receptor

In reviewing the microarray dataset, we were surprised to find that *Adra1a* was the sixth most downregulated transcript (**Table 1**). Our lab has been studying *Adra1a*, the gene that encodes the α1A adrenergic receptor (α1A-AR) for over a decade but had never focused on its transcriptional regulation, of which relatively little is known. To determine whether ROR⍺ regulates *Adra1a* expression, we examined transcript levels in multiple experimental models using quantitative reverse transcription PCR (qRT-PCR). In the global loss-of-function context, *Adra1a* expression was 3.1-fold lower in ROR⍺^sg/sg^ (n=6) than WT (n=12) hearts (p=0.0003, **Figure 2A**). To ascertain whether this relationship was cardiomyocyte-autonomous or might arise from systemic loss of RORα function, we assayed transcript abundance in the cardiomyocyte-specific RORα knockout mouse (*Myh6*-Cre;*Rora*^fl/fl^ or CMKO) our lab created [15]. We found that CMKO hearts contained 2.1-fold less *Rora* mRNA than WT littermate hearts (n=8 per group, p<0.0001, **Figure 2B**).

**Figure 2.**
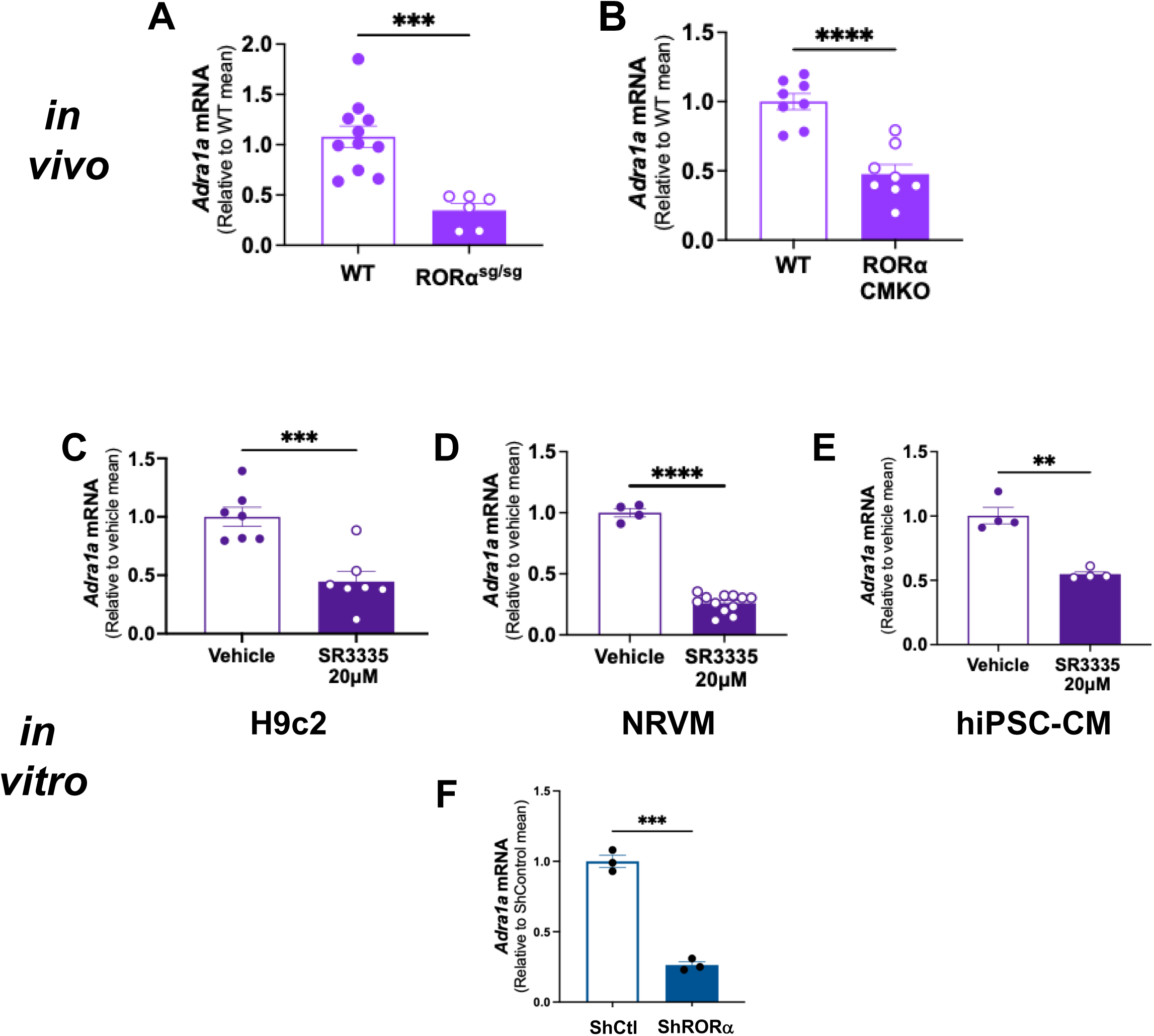
Genetic or pharmacologic loss of ROR⍺ function downregulates *Adra1a* mRNA. Abundance of *Adra1a* mRNA was quantified relative to *Actb* and *Tbp1* in **(A)** Wild-type (WT) and ROR⍺^sg/sg^ mouse hearts; (**B**) WT and cardiomyocyte-specific RORα knockout (CMKO) mouse hearts. *In vitro* experiments measured *Adra1a* mRNA after 24h exposure to either the RORα antagonist SR3335 or vehicle in (**C**) H9c2 ventricular myoblasts, **(D)** Neonatal rat ventricular myoctyes (NRVMs) or (**E**) Human induced pluripotent stem cell derived cardiomyocytes (hiPSC-CMs); **(F)** *Adra1a* mRNA was compared in NRVMs infected with either lentiviral control shRNA or shRNA against *Rora*. Statistical significance was determined by Welch’s unpaired T-test. *p<0.05, **p<0.01, ***p<0.001, ***p<0.0001

**Table 1.** Top 7 Up- and Down-regulated transcripts from the RNA microarray.

| Top 7 Named Transcripts – Upregulated |  |  |  |  |
| --- | --- | --- | --- | --- |
| Gene | Change | Adj p-value | WT mean | sg/sg mean |
| Gprin1 | 15.2 | 0.010 | 8.1 | 91.8 |
| Cacng3 | 15.5 | 0.043 | 192.6 | 2735.7 |
| Akap7 | 18.3 | 0.030 | 46.7 | 1457.8 |
| Cpa2 | 19.0 | 0.019 | 154.1 | 2260.1 |
| Cd247 | 19.2 | 0.028 | 2.4 | 130.4 |
| Clec2e | 19.3 | 0.001 | 3.1 | 57.9 |
| Adamts7 | 22.8 | 0.002 | 305.9 | 6453.4 |

**Table 1. Top 7 Up- and Down-regulated transcripts from the RNA microarray**
| Top 7 Named Transcripts – Downregulated |  |  |  |  |
| --- | --- | --- | --- | --- |
| Gene | Change | Adj p-value | WT mean | sg/sg mean |
| Mlec | -240.3 | 0.008 | 1935.2 | 32.3 |
| Etf1 | -128.4 | 0.006 | 1019.6 | 16.2 |
| Cpsf2 | -106.1 | 0.001 | 1100.7 | 10.2 |
| Atf1 | -102.6 | 0.005 | 1998.8 | 34.1 |
| Creg1 | -100.7 | 0.022 | 1433.4 | 88.8 |
| Adra1a | -99.1 | 0.019 | 1887.1 | 84.4 |
| Eif2a | -98.4 | 0.006 | 3831.3 | 74.1 |

We then pursued pharmacologic loss-of-function approaches *in vitro* to complement our genetic loss-of-function *in vivo* findings. We incubated H9c2 ventricular myoblasts, neonatal rat ventricular myocytes (NRVMs) or human iPSC-derived cardiomyocytes (hiPSC-CMs) with the ROR⍺ inverse agonist SR3335[22] or vehicle for 24 hours. We found that blocking ROR⍺ activation resulted in 2.3-fold (p=0.0006) *Adra1a* downregulation in H9c2s (**Figure 2C**), 3.8-fold (p<0.0001) downregulation in NRVMs (**Figure 2D**), and 1.8-fold downregulation (p=0.003) in hiPSC-CMs (**Figure 2E**) relative to vehicle controls.

Lastly we used a genetic loss-of-function with a validated lentiviral shRNA against *Rora*[14, 15] in NRVMs and identified a 3-fold downregulation of *Adra1a* compared to NRVMs infected with a scrambled control shRNA (**Figure 2F**, p = 0.0005).

In summary, we used an unbiased transcriptomic approach in a global loss-of-function mouse model to identify a novel putative regulation of α1A-ARs by ROR⍺ in the heart. We then employed directed qRT-PCR assays to confirm this finding *in vivo*. Our *in vitro* studies, using pharmacologic loss-of-function in multiple cardiac cell types, corroborated our *in vivo* studies. Collectively, these findings identified ROR⍺ as a potential transcriptional regulator of *Adra1a* expression in cardiomyocytes. The consistent findings in mouse, rat and human models suggest that this relationship is conserved across species.

### RORα gain-of-function enhances *Adra1* expression in cardiomyocytes

We then carried out pharmacologic and genetic gain-of-function experiments to complement our loss-of-function approaches. To evaluate whether RORα activation is sufficient to drive *Adra1a* expression, we first cultured H9c2s, NRVMs, and hiPSC-CMs in the RORα agonist SR1078 [23, 24] for 24 hours. We found that activation of RORα led to 2.6-fold induction (p<0.0001) of *Adra1a* in H9c2s (**Figure 3A**), 2.6-fold (p<0.0001) induction in NRVMs (**Figure 3B**), and 1.9-fold (p=0.03) induction in hiPSC-CMs (**Fig. 3C**).

**Figure 3.**
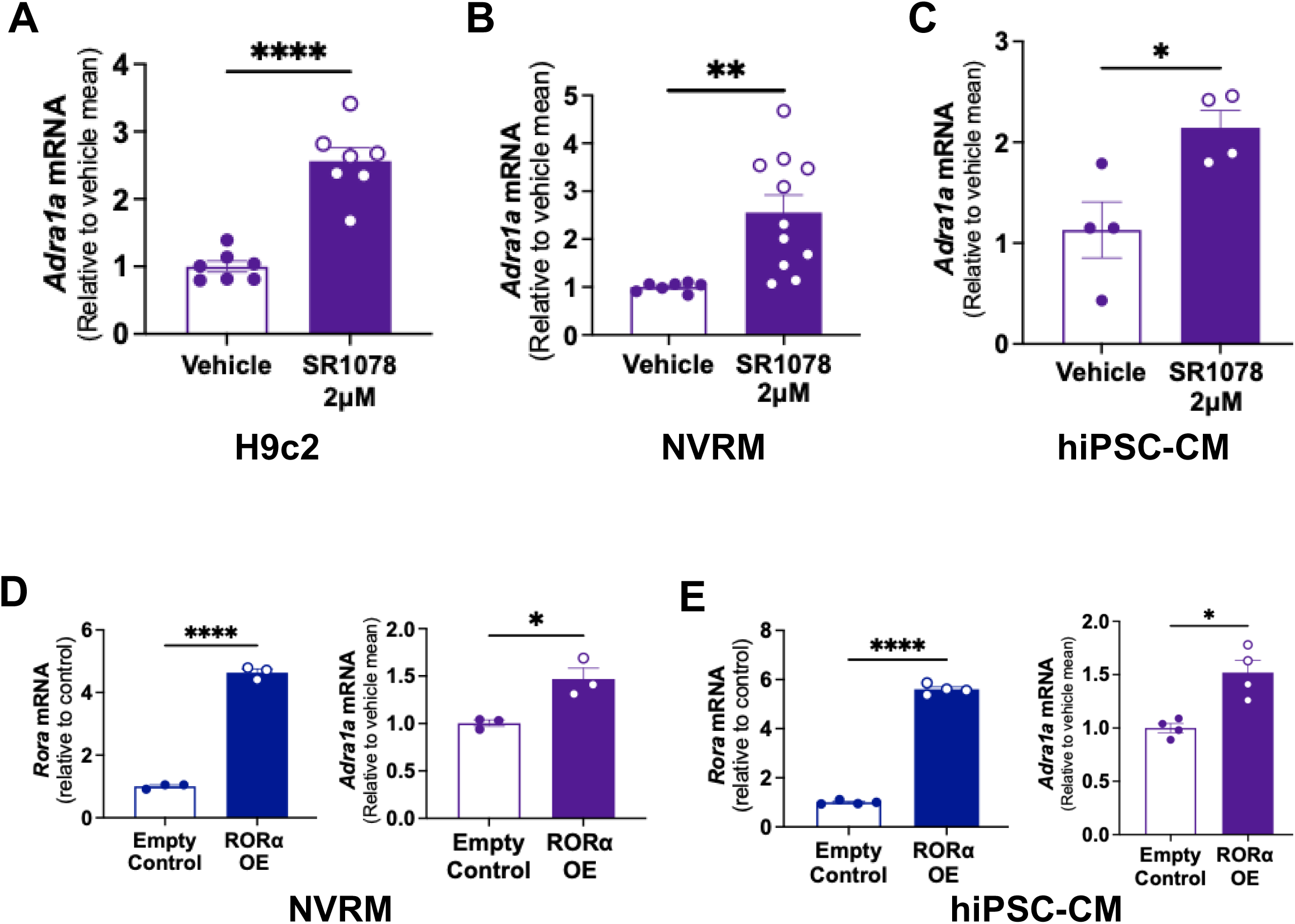
Genetic or pharmacologic gain of RORá function promotes *Adra1a* expression in cardiomyocytes. *Adra1a* mRNA was measured after 24h exposure to either the RORα agonist SR1078 or vehicle in (**A**) H9c2 ventricular myoblasts, **(B)** Neonatal rat ventricular myoctyes (NRVMs) or (**C**) Human induced pluripotent stem cell derived cardiomyocytes (hiPSC-CMs) or after lentiviral-mediated over-expression of RORα (72 h, 10 MOI) achieved in **(D)** NRVMs or (**E**) hiPSC-CMs with empty lentivirus infection as control. n ≥ 3 independent experiments. *p<0.05, **p<0.01, ***p<0.001, ***p<0.0001.

Next, we utilized a lentiviral construct that drives RORα overexpression (RORα-OE) for a genetic gain-of-function approach. We infected NRVMs and hiPSC-CMs with RORα-OE (MOI 10) or empty vector lentiviral control then carried out qRT-PCR for *Adra1a* after 72 hours. We found that the RORα-OE lentivirus induced 4.6-fold (p<0.0001) overexpression of RORα in NRVMs, resulting in a 1.4-fold (p=0.02) induction of *Adra1a* (**Figure 3D**). In hiPSC-CMs, RORα-OE infection induced 5.6-fold (p<0.0001) overexpression of RORα, leading to 1.5-fold (p=0.02) induction of *Adra1a* (**Figure 3E**). The relatively modest increase in *Adra1a* abundance may reflect the fact that endogenous RORα is already highly expressed and bound to *Adra1a*, though not fully activated by agonist.

Collectively these findings establish that both pharmacologic and genetic gain of RORα function is sufficient to enhance *Adra1a* expression in rodent and human cardiomyocytes, further suggesting that RORα is a positive transcriptional regulator of α1A-ARs in the heart.

### RORα enhances hypoxia-induced *Adra1a* upregulation

It has long been recognized that the sympathetic nervous system is activated robustly in the setting of myocardial ischemia (reviewed in [25, 26] and others). As part of that critical response, α1-AR expression is upregulated 2-fold, whereas β1-AR expression increases only roughly 30%[26]. The transcriptional regulators of this hypoxia-induced α1-AR induction are unknown. We[15] and others[27, 28] have demonstrated that RORα is upregulated in cardiovascular tissues in the setting of hypoxia.

To investigate whether RORα contributes to the upregulation of *Adra1a* expression under hypoxic conditions, we housed WT and RORα-CMKO mice at 10% oxygen tension in a hypoxia chamber for 8 hours then immediately harvested heart tissue. In WT mice, *Adra1a* expression increased 1.6-fold. In CMKO mice, *Adra1a* levels were lower in normoxia as expected (see **Figure 2**), and there was no induction of *Adra1a* under hypoxic conditions. (**Figure 4A**).

**Figure 4.**
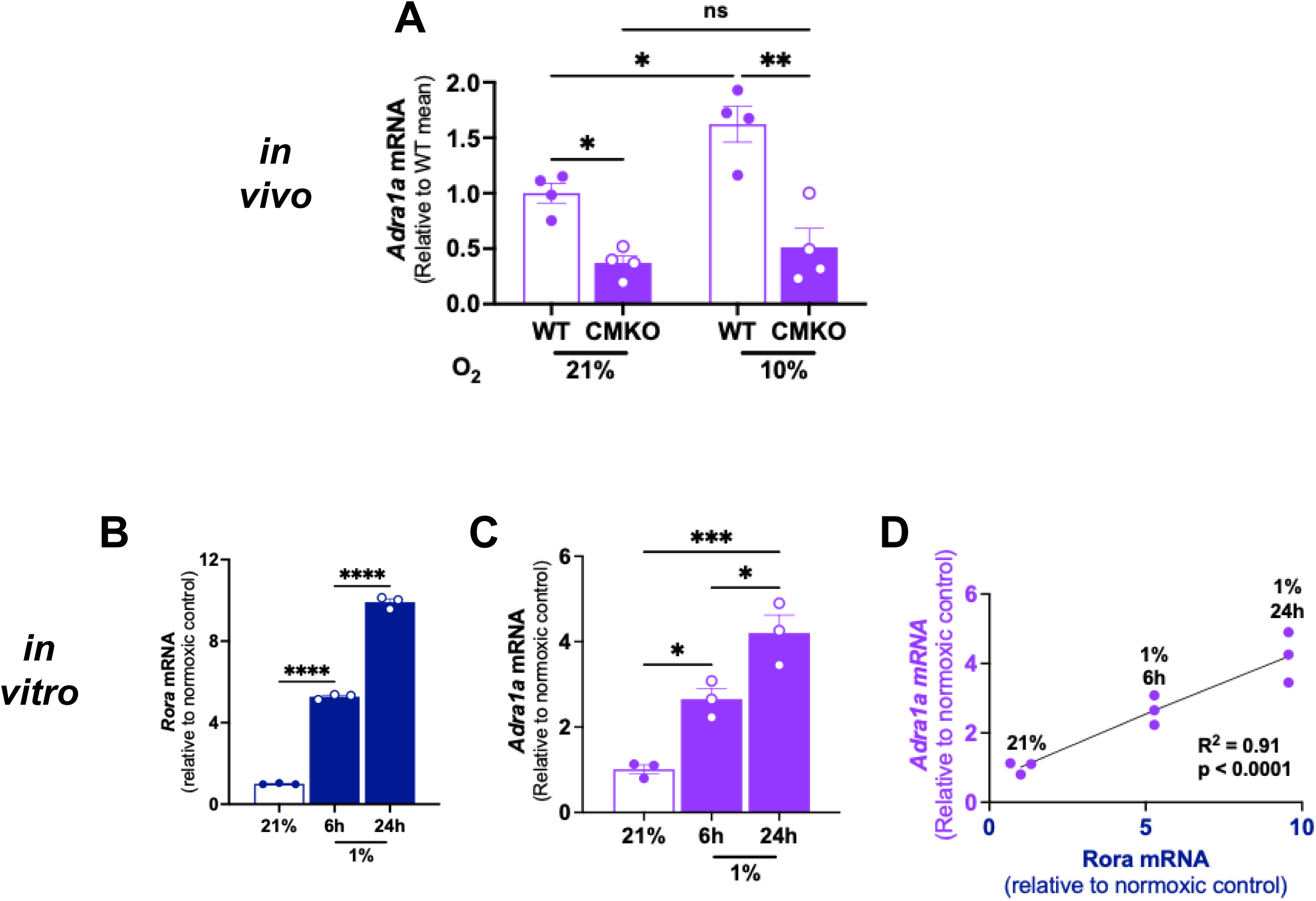
RORá is essential for hypoxia-mediated *Adra1a* upregulation. Quantitative RT-PCR under normoxia or hypoxia (10% *in vivo* or 1% *in vitro*) compared (**A**) *Adra1a* mRNA in cardiomyocyte-specific RORα knockout (CMKO) mice; (**B**) *Rora* mRNA or **(C)** *Adra1a* in human induced pluripotent stem cell derived cardiomyocytes (hiPSC-CMs). **(D)** Linear regression analysis of *Rora* and *Adra1a* in hiPSC-CMs *p<0.05, **p<0.01, ***p<0.001, ***p<0.0001 by two-way ANOVA (A) or one-way ANOVA with Tukey’s post-hoc analysis (B and C)

To extend these findings to a cardiomyocyte-autonomous *in vitro* context, we exposed hiPSC-CMs to low oxygen tension (1% O₂) for 6 or 24 hours. Hypoxia induced a 5.3-fold upregulation of *Rora* at 6h and a 9.9-fold upregulation at 24h (p<0.0001 vs normoxia for both timepoints, **Figure 4B**). *Adra1a* transcript abundance increased 2.7-fold at 6h (p=0.02) and 4.2-fold at 24h (p=0.0005) at 24h (**Figure 4C**). We then applied linear regression analyses to these findings and identified a strong correlation between *Rora* and *Adra1a* abundance (R^2^=0.91, p<0.0001, **Figure 4D**)

Taken together, these findings suggest that RORα contributes to the upregulation of α1A-ARs under hypoxic conditions, a central aspect of the sympathetic nervous system response to ischemia in the heart.

### RORα transcriptionally regulates the human *ADRA1A* promoter

Given that RORα functions primarily as a transcription factor, we reasoned that the correlations that we observed between RORα activity and *Adra1a* abundance most likely arose from direct transcriptional regulation of *Adra1a* by RORα. We previously have shown that RORα transactivates IL6[14] and Caveolin-3[15] by binding to ROREs present in the proximal promoter, but no previous studies have demonstrated that RORα regulates *Adra1a* transcription. To investigate whether RORα similarly regulates the *ADRA1A* promoter, we cloned an ∼819 bp proximal promoter fragment upstream of the human *ADRA1A* gene into the pGL4.10 luciferase reporter vector. *In silico* analysis identified two RORE motifs located at positions –721 to –727 (E2) and –624 to –630 (E1).

To assess whether RORα binds to these motifs, we generated a mutant construct in which the E2 site in the *ADRA1A* promoter was altered from TGGGTCA to TG<u>A</u>G<u>C</u>CA (**Figure 5A**). Luciferase reporter assays revealed that RORα overexpression in H9c2 ventricular myoblasts increased *ADRA1A* promoter activity in concentration-dependent manner (**Figure 5B**). Co-treatment with the RORα agonist SR1078 further enhanced luciferase activity, whereas the inverse agonist SR3335 completely abolished promoter activation **(Figure 5B)**.

**Figure 5.**
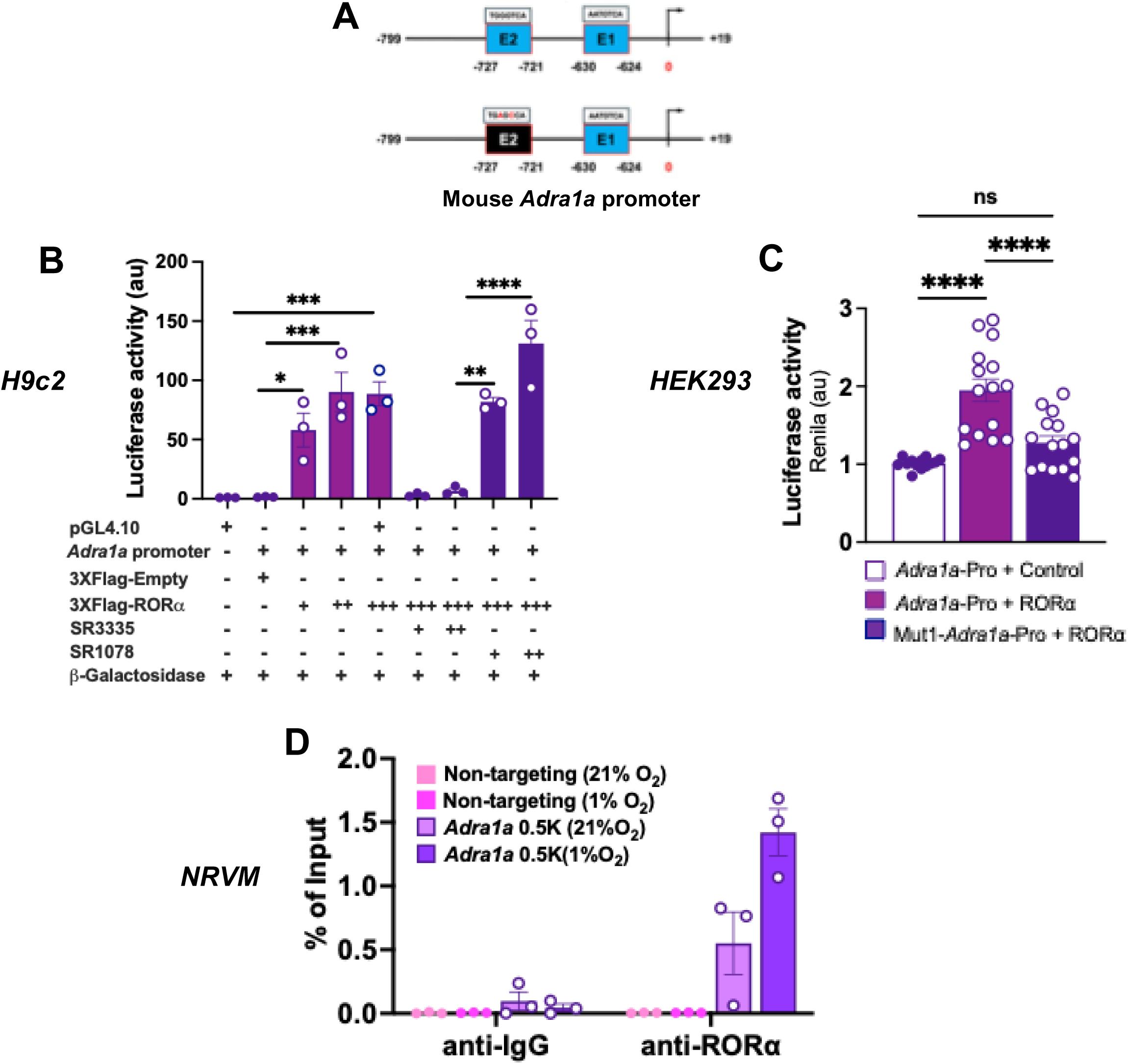
Activation of the mouse *Adra1a* promoter by RORα. **(A)** Schematic representation of proximal promoter region of mouse *Adra1a* consisting E-box 1 and 2 and mutant E-box 2. **(B)** Luciferase construct encompassing mouse *Adra1a* promoter (pGL4.10-ADRA1α-Pro) transfected into NIH3T3 fibroblasts with a RORα expression vector (p3xFLAG-CMV-RORα) or empty vector control (p3xFLAG-CMV), followed by treatment either with RORα agonist SR1078 or inverse agonist SR3335, with normalization to co-transfected with β-galactosidase construct for luciferase activity. **(C)** pGL4.10-Adra1a-Pro or mutant E-Box 2 of human *ADRA1A* promoter (Mut1-pGL4.10-*ADRA1a*-Pro) transfected into HEK293 cells with p3xFLAG-CMV-RORα or p3xFLAG-CMV and co-transfected with Renilla luciferase pRL for normalization (**D**) Chromatin immunoprecipitation (ChIP)-PCR of *Adra1a* in NRVMs exposed to normoxic or hypoxic conditions exposed NRVMs incubated with anti-RORα or anti-IgG (control) antibodies. Statistical significance was determined by one-way ANOVA with Tukey’s post-hoc analysis (B and C) or two-way ANOVA with Tukey’s post-hoc analysis (D). *p<0.05, **p<0.01, ***p<0.001, ***p<0.0001

We then cloned our mutant RORE(2) construct into HEK293 cells and found that mutation of the RORE2 site markedly reduced luciferase activity, confirming that RORα-mediated transactivation requires an intact RORE(2) of the *ADRA1A* promoter (**Figure 5C**). To further validate the finding that RORα binds and regulates the *Adra1a* promoter in cardiomyocytes, we performed chromatin immunoprecipitation PCR(ChIP-PCR), on NRVMs cultured in normoxic or hypoxic conditions. We found that an anti-RORα antibody, but not control anti-IgG, bound RORE(2) in the *Adra1a* and that hypoxia further enhanced binding (**Figure 5D**), consistent with our *in vivo* and *in vitro* findings (**Figure 4**).

Collectively, these results demonstrate that RORα directly transactivates the *ADRA1A* promoter via binding to the proximal RORE(2) motif. Pharmacological activation and hypoxia further enhance this transcriptional regulation.

### Activation of RORα enhances α1A-AR-mediated ERK signaling

We then sought to determine whether the transcriptional regulation of *Adra1a* by RORα was accompanied by an increase in functional α1A-ARs. As there are no selective antibodies for α1A-ARs[29], we used the well-recognized cardioprotective activation of ERK1/2 by α1A-ARs[30–32] as a readout. We cultured NRVMs in the presence or absence of the selective α1A-AR agonist A61603 (10nM) and/or the RORα agonist SR1078 (2μM) then immunoblotted for phosphorylated (p-ERK), total ERK (t-ERK) and the loading control GAPDH (**Figure 6A**). SR1078 alone did not affect ERK activation but A61603 increased p-ERK abundance 11.0-fold relative to GAPDH, and the addition of SR-1078 led to 27.1-fold induction of ERK phosphorylation (**Figure 6B**). Relative to t-ERK, A61603 and A61603 with SR1078 increased ERK activation 13.3-fold and 32.5-fold respectively (**Figure 6C**). These findings demonstrate an important functional consequence to the transcriptional regulation of *Adra1a* by RORα.

**Figure 6.**
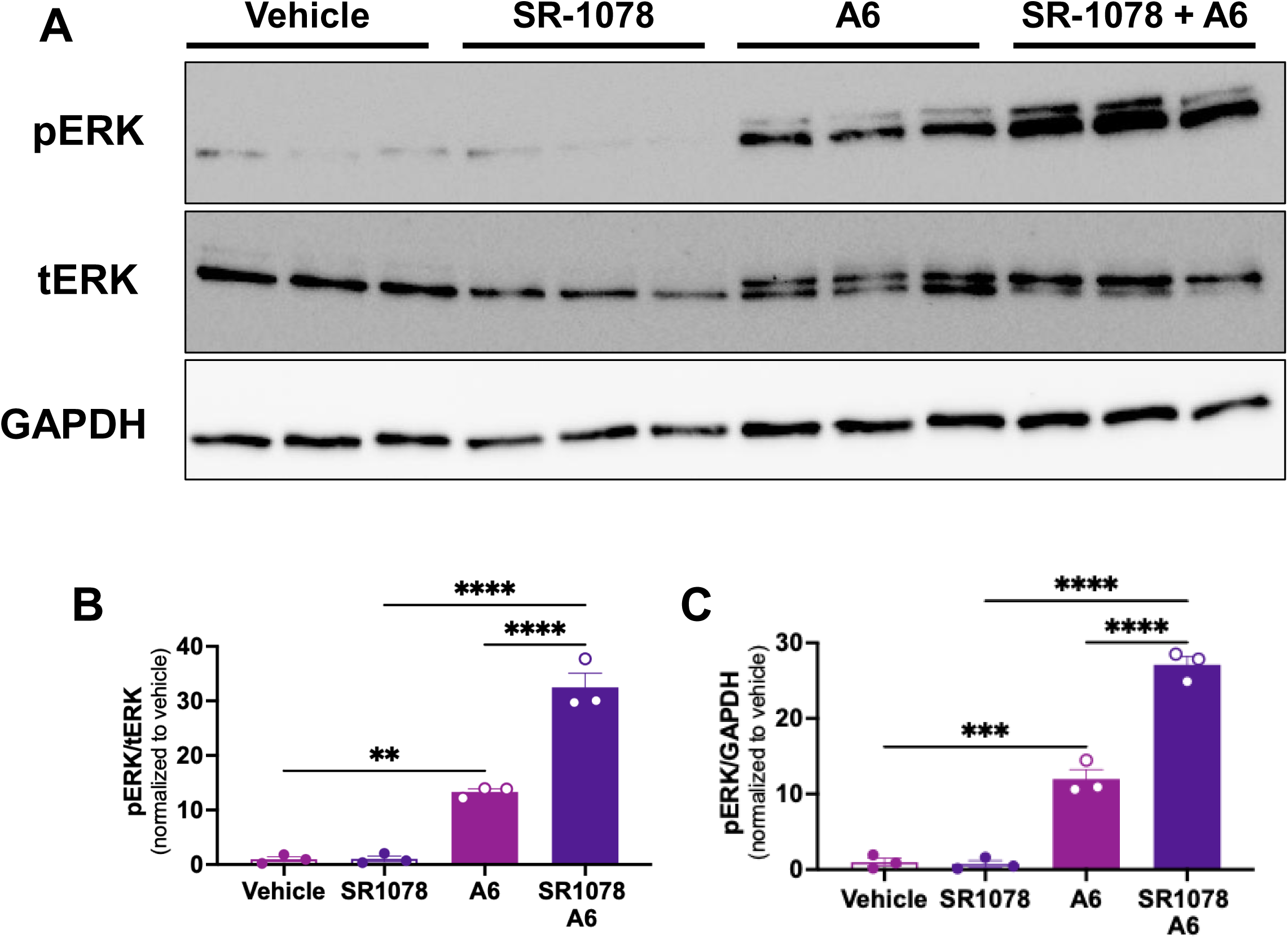
RORα agonist augments α1A-AR mediated ERK signaling. NRVMs were exposed to RORα agonist SR1078 (2μM) and/or α1A-AR agonist A61603 (10nM) for 24h (**A**) Representative immunoblot; Summary densitometry for pERK1/2 relative to **(B)** total ERK1/2 or (**C**) GAPDH. Statistical significance was determined by one-way ANOVA with Tukey’s post-hoc analysis. *p<0.05, **p<0.01, ***p<0.001, ***p<0.0001.

### RORα oscillates with the same periodicity as *Adra1a*

RORα is a well-established regulator of circadian biology[33], integrating clock control with metabolic and cellular signaling outputs (reviewed in [6]). Unsurprisingly, our RNA microarray analysis identified broad differences in the expression of transcripts in the GO term Circadian Entrainment (**Figure 7A**). Interestingly, the CircaDB database has identified *Adra1a* as the only adrenergic receptor that oscillates in the heart[34]. To test whether RORα might modulate circadian oscillation of *Adra1a*, we harvested the hearts of C57B6N mice every at 4-hour intervals, using qRT-PCR to assay for the abundance of *Rora* and *Adra1a* as well as the positive oscillatory control *Bmal1 (Arntl). Myh6* was assayed as a non-oscillatory negative control. We found that *Rora* and *Adra1a* transcripts oscillated with the same periodicity. *Bmal1* oscillated with alternate periodicity and *Myh6* exhibited no circadian variation in abundance (**Figure 7B**). Though published studies using other tissues have demonstrated that *Rora* directly positively regulates *Bmal1* transcription*[33]*, CircaDB reveals that they oscillate with staggered periodicity in the heart, very similar to our findings. In future studies, we will be very interested in pursuing the physiological implications of RORα-mediated circadian regulation of α1A-AR expression in the heart.

**Figure 7.**
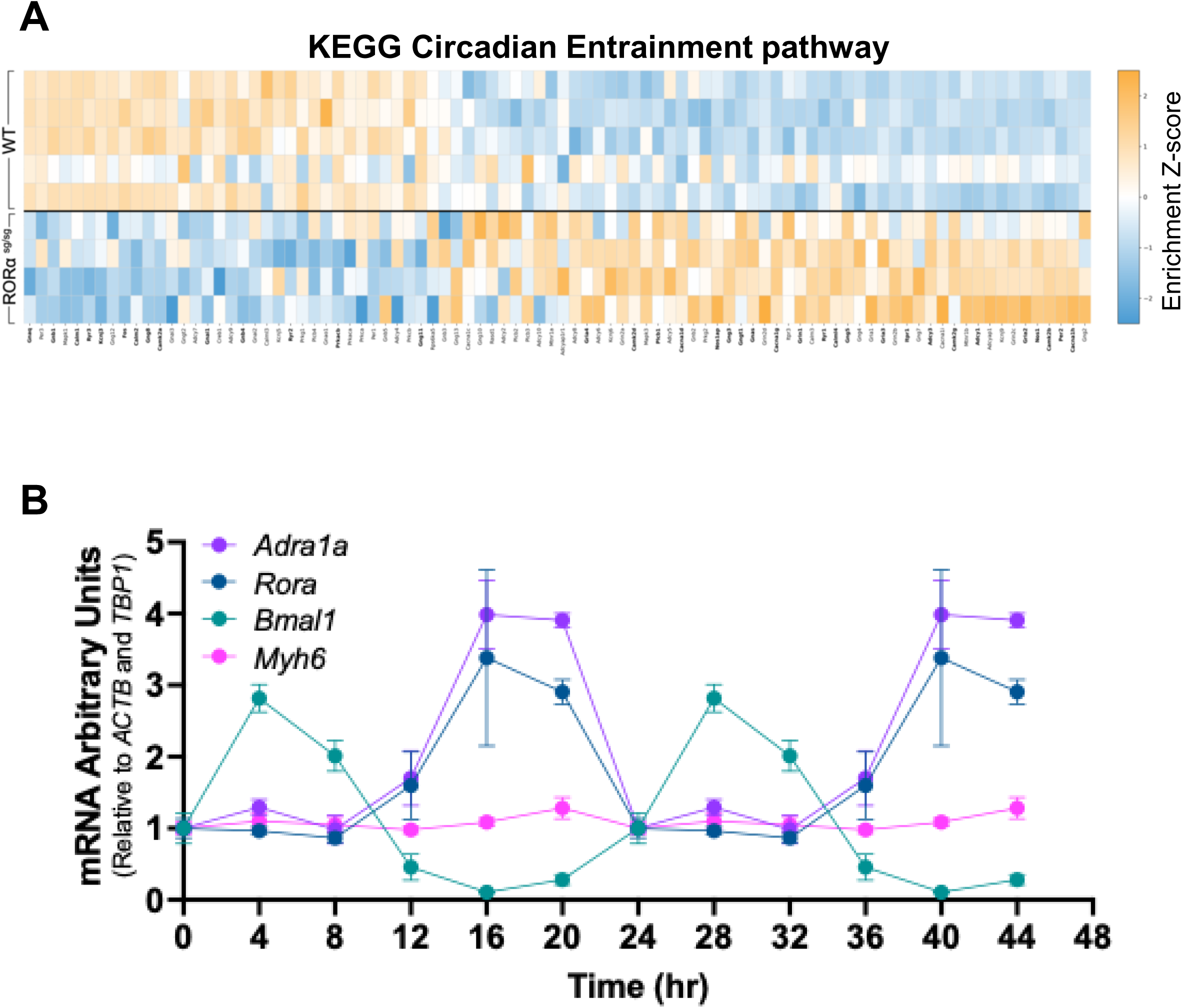
*Rora* oscillates with the same periodicity as *Adra1a*. **(A)** KEGG circadian entrainment pathway analysis of RNA microarray data. **(B)** qRT-PCR analysis of mRNA expression in wild type mouse hearts collected at synchronized time points. Data are shown as mean ± SEM from three independent experiments.

## Discussion

In this manuscript we show that the absence of functional RORα confers broad transcriptomic effects on the heart that likely collectively result in the atrophic and hypocontractile cardiac phenotype that we previously published for the RORα^sg/sg^ mice. We also demonstrate that RORα directly regulates transcription of *Adra1a*, the gene that encodes the α1A subtype of α1-ARs. Collectively these novel findings position RORα as another nuclear receptor with important regulatory roles in cardiac structure and function, including myogenesis and transduction of sympathetic nervous system activation.

Our microarray data identify a decrease in the expression of many sarcomeric genes suggesting impaired myogenesis. As a functional consequence of the relative dearth of contractile proteins, RORα^sg/sg^ mice have reduced fractional shortening compared to WT littermates[14]. The microarrays also reveal reduced expression of multiple critical transcriptional regulators of myogenesis, including TEAD-1 (TEF-1)[35], MEF2a[36], GATA4[37], and Srf[38]. Each of these transcription factors has at least one ROR response element (RORE), the consensus DNA binding motif for RORα (AGGTCA preceded by a 6bp A/T rich region) in its promoter region, suggesting that they might well be RORα target genes. As such, it is possible that RORα regulates myofibrillogenesis at multiple levels: directly as a co-regulator of sarcomeric proteins and indirectly as a regulator of myogenic transcription factor abundance. Though a role for RORα in myofibrillogenesis and hypertrophy in cardiac muscle has not been previously described, RORα mediates differentiation of C2C12 myotubes into skeletal muscle cells through interaction with p300 and MyoD[39]. Our findings position RORα as the third known nuclear receptor to participate directly in transcriptional regulation of sarcomeric and calcium-handling genes, joining estrogen related receptors alpha and gamma (ERRα and γ)[40].

Published studies from other tissues indicate that RORα functions primarily as a ligand dependent transcription factor. Metabolites of cholesterol, vitamin D and the omega-3 fatty acid docosahexaenoic acid (DHA) bind reversibly to the RORα ligand binding domain (LBD)[41, 42] to enhance or repress transcriptional activation of genes containing an RORE in the promoter of target genes[5, 43]. RORα also interacts with multiple co-repressors and co-activators, including PGC1α, p300, and CBP, to regulate transcription in a ligand-independent manner[6]. It is unclear whether RORα exerts its effects in the heart through ligand-dependent or independent activity and the endogenous ligand for cardiomyocyte RORα remains unknown. Contrary to some reports[7, 44] melatonin is not a bona fide ligand for RORα[45].

From the many differentially expressed transcripts with well-recognized critical roles in cardiac biology, we chose to focus on the regulation of *Adra1a (*α1A-AR) expression given our longstanding interest in this receptor. We and others have demonstrated that α1-ARs play protective roles in the heart that contrast with and mitigate the toxic effects of chronic β-AR activation in the failing heart[46–48]. More recently we have shown that these adaptive functions are mediated by the α1A-AR subtype[49] and that α1A-AR activation protects against anthracycline-induced cardiotoxicity[50] and myocardial infarction[51]. We also recently showed that α1A-selective antagonists, including the widely prescribed medication tamsulosin (Flomax), are associated with an increased risk of all-cause mortality within one year of the initial prescription[52].

Relatively little is known about the transcriptional regulation of α1A-ARs. One study in neuroblastoma cell line found that Sp1 binds GC boxes in the *Adra1a* promoter and drives basal transcriptional activity[53]. Functional promoter analyses in rat cardiomyocytes identified a hypoxia-responsive element and additional GATA and CREB motifs though did not confirm direct promoter binding and regulation[54] and KLF15 has been shown to directly bind enhancers that stimulate *Adra1a* expression[55]. To our knowledge, however, ours is the first study to demonstrate direct transcriptional regulation of cardiomyocyte *Adra1a*.

On the other hand, it has long been recognized that activation of α1-ARs results in enhanced transcription of a number of sarcomeric genes, mediated by increased activity of critical myogenic transcription factors including TEF1/TEAD[56, 57], GATA4[58, 59], and CREB[60]. These earlier publications, coupled with our new demonstration that RORα enhances *Adra1a* transcription, raises the possibility that RORα participates in a feed-forward transcriptional program that transduces some effects of sympathetic nervous system activation on the cardiomyocyte myofibrillar apparatus.

The primary limitation of our RNA microarray findings is the use of a mouse model that globally lacks functional RORα, leading to metabolic derangements[6] that could affect the heart. As such, we cannot exclude a contribution from extracardiac RORα in our experiments that utilized the RORα^sg/sg^ mice (**Figures 1 and 2A**). However, we also find in focused studies that cardiomyocyte-specific RORα knockout mice have lower levels of *Adra1a* than WT littermates (**Figures 2B and 4A**). Additionally, our *in vitro* experiments using three distinct cardiomyocyte platforms (**Figures 2C-E, 3, 4, 5 and 6**) strongly suggest direct and cell autonomous roles for RORα in regulating *Adra1a* transcription.

We long have advanced the concept that selective α1A-AR agonists represent a promising novel approach to the treatment of HF[46, 49]. Interestingly, recent studies of non-selective RORα agonists have shown potential cardioprotective effects. Maresin-1, an endogenous RORα ligand, promotes physiological cardiomyocyte hypertrophy through IGF1[61]. The naturally occurring flavonoid small molecule nobiletin protects against development of metabolic syndrome in mice fed a high fat diet[62]. However, selective synthetic agonists of RORα have not been tested in the context of heart injury, though SR1078 was well tolerated and reduced repetitive behaviors in an animal model of autism[23]. Any potential contribution of enhanced α1A-AR activity to these promising findings has not been explored.

