## Supplemental Figure for "The nuclear receptor ROR-alpha is a critical myogenic regulator of the cardiomyocyte transcriptome, including the alpha-1A adrenergic receptor"

### Supplemental Figure S1

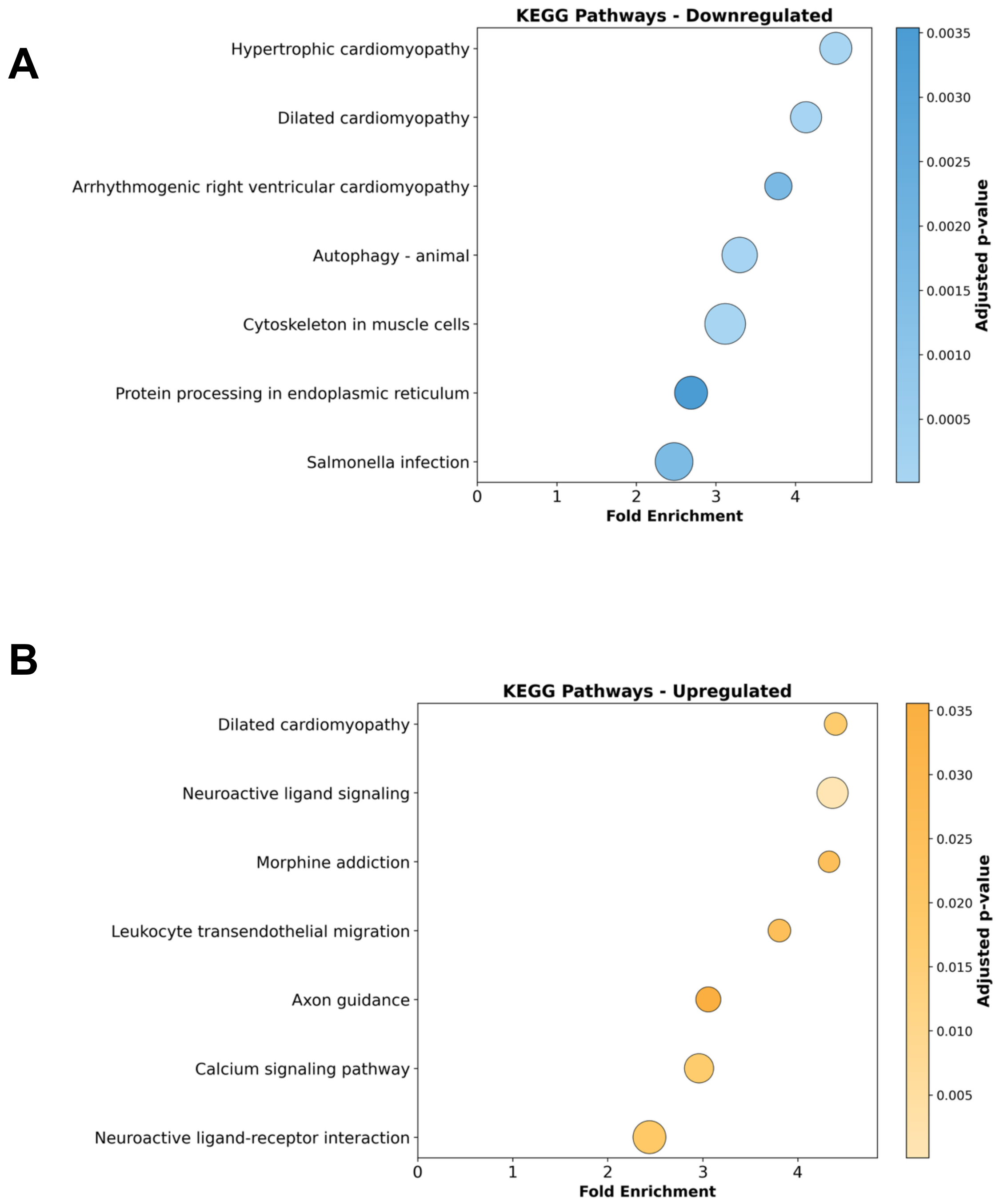


**Supplemental Figure S1.** KEGG pathway analysis of **(A)** 2-fold downregulated and **(B)** 2-fold upregulated transcripts in RORα$^{sg/sg}$ (n=4) vs WT (n=5) hearts.
